# Multiple Oscillatory Neural Rhythms Support Metacognitive Access of Working Memory

**DOI:** 10.64898/2026.04.08.717328

**Authors:** Yang Di, Xingwei An, Hsin-Hung Li

**Affiliations:** Academy of Medical Engineering and Translational Medicine, Tianjin University, Tianjin, 300072, China; Department of Psychology, The Ohio State University, Columbus, OH 43201, USA

**Keywords:** working memory, metacognition, EEG, uncertainty, confidence, oscillation

## Abstract

Working memory enables the maintenance of information to guide behavior, yet its representations are inherently noisy and variable. To act effectively, observers must track the uncertainty of their own memory. However, the neural mechanisms supporting uncertainty representations in working memory remain poorly understood, particularly at the level of neural dynamics. Here, we combined electroencephalography (EEG) with a spatial visual working memory task that elicited trial-by-trial uncertainty reports to investigate how oscillatory activity encodes memory uncertainty. We identified two forms of uncertainty supported by distinct oscillatory neural activity. Using a probabilistic encoding–decoding model, we found that memory content can be reconstructed from alpha-band activity, and that the precision of these representations predicts subsequent uncertainty reports, consistent with probabilistic mnemonic representations. In parallel, beta-band activity contained a scalar, magnitude-based, representation of uncertainty. This signal exhibited slow, persistent dynamics across task epochs and tracked uncertainty reports for both the current and previous trials. Together, these findings show that the human brain multiplexes two neural codes for memory uncertainty, representational precision and a scalar confidence signal, across distinct oscillatory rhythms, providing a neural dynamical account of metacognitive access to working memory.

## Introduction

Working memory enables neural representations to be maintained beyond the moment of perception and to guide behavior. A hallmark of working memory is that its quality fluctuates over time (Fougnie et al. 2012; van den Berg et al. 2012; Bays et al. 2024). Even when identical stimuli are encoded repeatedly, the neural activity supporting mnemonic representations remains variable and noisy, rendering working memory uncertain. Adaptive behavior therefore requires access not only to remembered content but also to its uncertainty, enabling individuals to evaluate how much their memory can be trusted. This metacognitive process is sometimes referred to as metamemory or meta-WM (Miyamoto et al. 2017; Ning et al. 2026; Baird et al. 2013; Hu et al. 2021).

While the neural basis of uncertainty and metacognitive judgments has been extensively studied in perception, (Walker et al. 2020; van Bergen et al. 2015; van Bergen and Jehee 2019; reviewed in Walker et al. 2023; Bang and Fleming 2018; Kepecs et al. 2008; Fleming et al. 2012; Kiani and Shadlen 2009; Vivar-Lazo and Fetsch 2026; Okazawa et al. 2021), only recently have studies begun to investigate how the human brain represents uncertainty in working memory. Recent human fMRI studies show that the quality of mnemonic representations in visual and intraparietal cortex correlate with behavioral uncertainty reports (Li et al. 2021, 2025). However, metacognition likely relies on multiple signals rather than solely on instantaneous memory fidelity. For example, uncertainty reports may be influenced by performance in recent trial history (Rahnev et al. 2015; Rouault et al. 2019). It remains unclear whether distinct formats of uncertainty are encoded by common neural representations or by multiple specialized codes. Moreover, the neural dynamics underlying the formation of memory uncertainty remain largely unexplored. In particular, it is unknown whether uncertainty is represented in oscillatory neural activity, a ubiquitous and temporally structured feature of population dynamics, despite its established role in working memory and higher-order cognition.

Oscillatory neural activity reflects coordinated population dynamics across frequency bands, each linked to distinct computations or cognitive functions (Fries, 2005; Buzsáki and Vöröslakos, 2023), making it well suited to support different aspects of metacognitive functions. Here we test whether and how two forms of uncertainty representation in working memory—representational precision and a scalar confidence signal—are carried by distinct oscillatory dynamics.

First, working memory uncertainty may arise from the precision of the mnemonic representations themselves. In this view, neural populations maintaining memory content encode not only a point estimate, but also probabilistic information whose precision reflects memory quality. In human scalp EEG (electroencephalogram) recordings, alpha-band oscillations (∼8–13 Hz) have been repeatedly implicated in visual working memory. During the memory delay, alpha activity is prominent and has been associated with spatial attention, suppression of irrelevant information, and maintenance of spatial representations (Spitzer and Blankenburg 2012; van Ede 2018; Jokisch and Jensen 2007; Samaha et al. 2016). Critically, the content of working memory can be reconstructed from alpha-band topography (Sutterer et al. 2019; Foster et al. 2017, 2016; Bae and Luck 2018; Bae 2021; Bender et al. 2025), indicating that alpha-band activity carries mnemonic content. However, prior studies rarely examined how these decoded memory content relate to subjective uncertainty or trial-by-trial memory error. Consequently, whether alpha-band activity supports uncertainty judgement of working memory remains unknown.

Second, metacognitive evaluation of working memory may rely on a different form of uncertainty representation. In addition to representational precision, neural populations may encode uncertainty as a simpler scalar signal that summarizes overall confidence level. Such signals could guide uncertainty reports or subsequent behavior without representing the remembered stimulus features themselves. Moreover, this signal may represent a confidence level that persists across trials or task epochs without interfering with ongoing sensory inputs or trialwise mnemonic representations.

In this study, we recorded electroencephalography (EEG) while human participants performed a spatial working memory task that required explicit trial-by-trial uncertainty reports. We found that uncertainty derived from decoded memory representations in alpha-band activity predicted subsequent subjective uncertainty judgments. In parallel, we identified a scalar uncertainty signal expressed most strongly in beta-band activity that tracked uncertainty across task epochs and across trials. Together, these findings show that working memory uncertainty is supported by multiple neural code across oscillatory dynamics, with representational precision and a scalar confidence signal carried by distinct frequency bands.

## Results

To explicitly probe working memory uncertainty that guides decisions, we adapted a paradigm that incentivized participants to report their memory uncertainty in a betting game (Honig et al. 2020; Li et al. 2021; Yoo et al. 2018). In a spatial visual working memory task, a brief target dot (500-msec, fixed at 10° eccentricity) was presented at a pseudo-random polar angle on each trial (Figure 1A). The target was followed by a 2-s delay period while participants maintained fixation at the central fixation. After the delay, participants first reported the remembered location using a mouse click. Immediately following this response, an arc representing a confidence interval was displayed on the screen, centered at the reported location. Participants adjusted the arc length to report their uncertainty about the remembered position. At the end of each trial, a feedback dot appeared at the true target location. Participants earned points if the target fell within the arc and earned zero points otherwise. When earned, the number of points decreased as a function of arc length. This task structure encouraged participants to report their memory uncertainty faithfully (Honig et al. 2020; Yoo et al. 2018).

**Figure 1.**
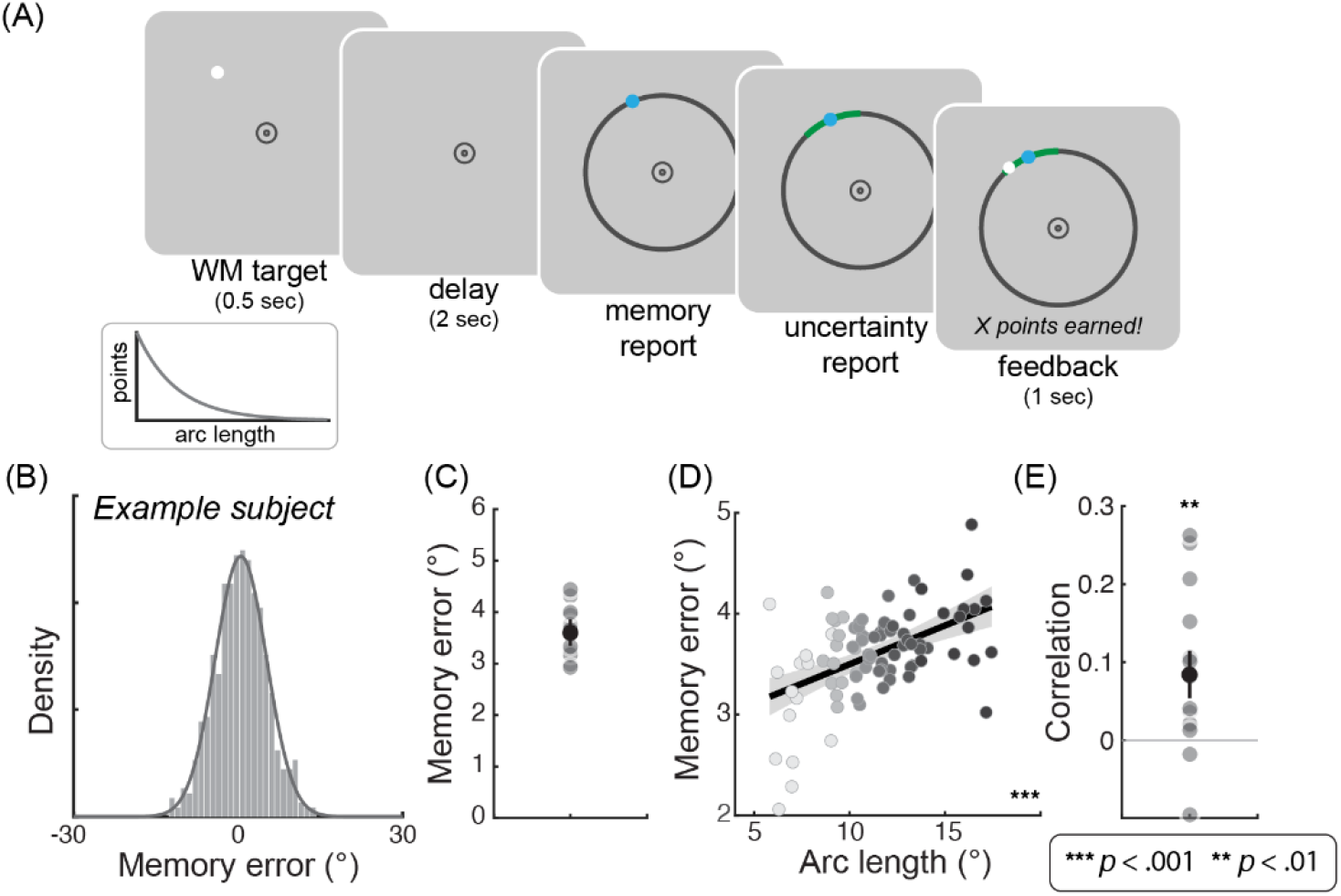
Procedures and behaviors. (A) Procedures. Each trial started with the target at a random polar angle followed by a delay period, during which participants maintained fixation and were required to remember the target location. After the delay, participants first reported their memory by clicking on the screen, and reported their uncertainty by adjusting the length of an arc. Feedback was subsequently provided as a dot at the true target position. If the target position fell within the arc, points were awarded accordingly based on the reward policy (lower left inset). (B) The distribution of behavioral memory error from an example participant. The gray line represents the best-fit von-Mises distribution. (C) Absolute behavioral memory error for individual participants (dots). The light gray dots represent group average. The black dot with the error bar represents group mean ±SEM. (D) Absolute memory error plotted against the arc length (reported uncertainty). Different gray levels indicate different bins sorted by arc length within each participant (8 bins per participant). The black line represents the best linear fit. (E) The correlation between memory error and arc length for individual participants (gray dots). The black dot represents the group average. The error bar represents ±SEM.

Participants’ memory reports were centered on the target location when expressed as memory error (Figure 1B). As expected given the presence of a single target and the relatively short delay duration, memory reports were quite precise (Figure 1C). Importantly, even within this narrow range of errors, participants demonstrated the ability to evaluate the quality of their memory. Reported arc length, an explicit subjective measure of uncertainty, increased with the magnitude of memory error (Figure 1D and 1E). These results are consistent with prior work (Honig et al. 2020; Li et al. 2021; Yoo et al. 2018), demonstrating that participants have metacognitive access to trial-by-trial fluctuations in working memory precision and that this task design effectively probes subjective uncertainty in working memory.

### Mnemonic representations in alpha-band activity reflect memory uncertainty

One potential source of memory uncertainty lies in the quality of the underlying mnemonic representations themselves. Rather than encoding only a single point estimate, memory representations may take a probabilistic format that inherently conveys uncertainty. We obtained a theoretically grounded neural measure of uncertainty by decoding working memory content from alpha-band activity with a generative model–based Bayesian approach (van Bergen and Jehee 2021; van Bergen et al. 2015; Li et al. 2021). In a cross-validation procedure, training data were used to construct a generative model that maps stimulus location (polar angle) to EEG activity patterns. The model assumed that multi-electrode EEG activity follows a multivariate normal distribution, with a mean that depends on the spatial tuning function of each electrode and a covariance that characterizes neural and measurement noise (see Methods). Once the model was established, we inferred target location on each single trial in the test set by computing a posterior distribution over locations given the observed EEG activity pattern *p*(*s* ∣b) through Bayesian inference (where *s* represents location and **b** represents alpha activity pattern; Figure 2A). Interpreting these decoded distributions as the mnemonic representations available to the observers, we used the posterior mean as a readout of remembered location and its width (standard deviation) as a neural index of memory uncertainty, which could guide the uncertainty report (Figure 2A).

**Figure 2.**
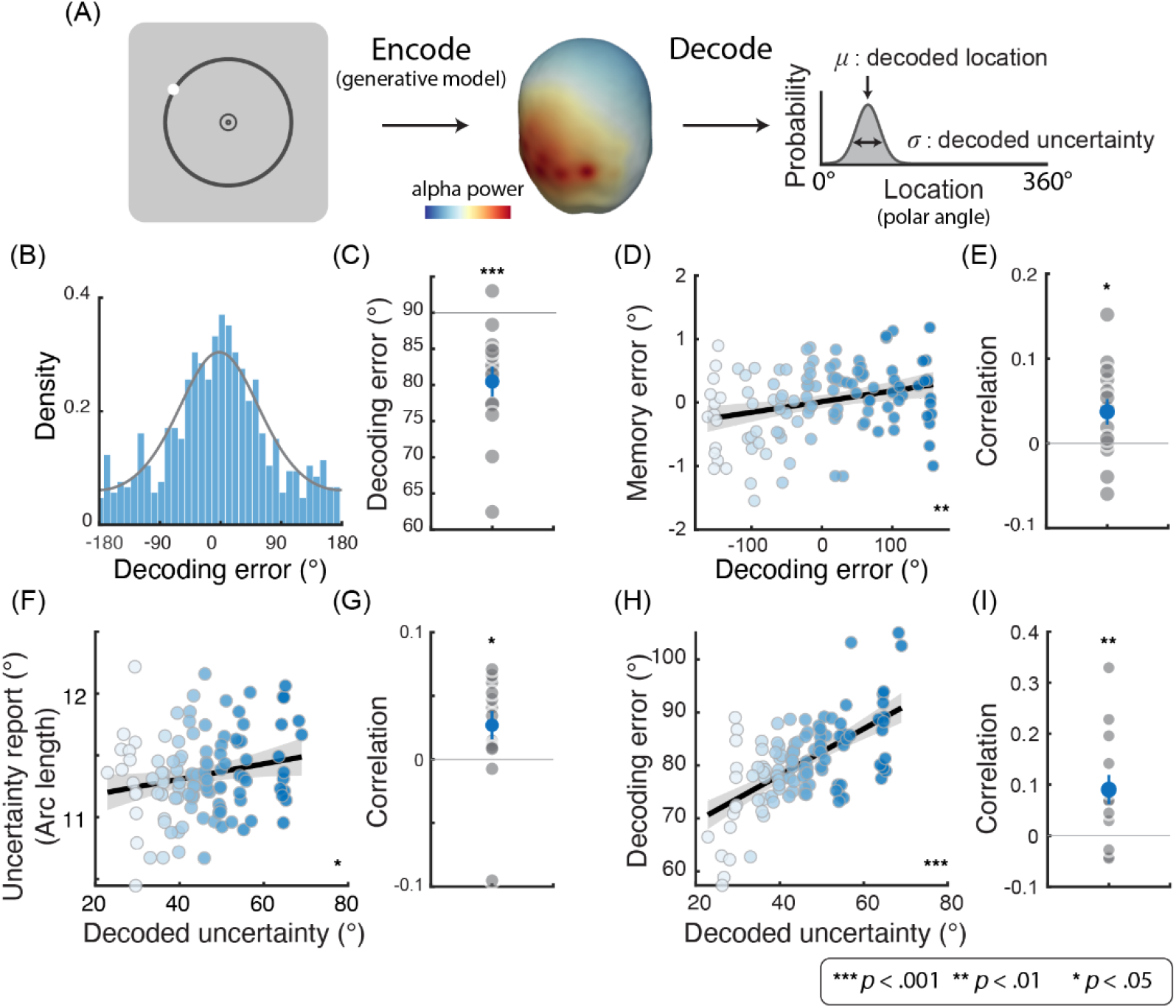
Probabilistic decoding of mnemonic representations from alpha-band activity. (A) Encoding model: We used the training data to build an encoding model, which modeled the multivariate alpha activity as a function of the polar angle of the target. Decoding: After fitting the encoding model, target location was inferred for each single trial in the test set using Bayesian inference. The posterior mean was taken as the decoded location, and the posterior standard deviation as the decoded uncertainty. (B) Decoding error histogram from an example participant. The gray linear represents the best-fit von Mises. (C) Absolute decoding error for individual participants (gray dots). The gray line represents the chance level (90°). The blue dot represents the group average and the error bar represents ±SEM. (D) Memory error plotted against decoding error. Different colors represent different bins sorted by decoding error within each participant (8 bins/colors per participant). The black line represents the best linear fit and gray shaded interval shows ±SEM. (E) Correlation between memory error and decoding error for individual participants (gray dots). The blue dot represents the group average. The error bar represents ±SEM. (F) Uncertainty report plotted against decoded uncertainty. Different colors indicate different bins sorted by decoded uncertainty within each participant. The black line represents the best linear fit. (G) Correlation between uncertainty report and decoded uncertainty for individual participants (gray dots). The blue dot represents the group average. The error bar represents ±SEM. (H) Absolute decoding error plotted against decoded uncertainty. Different colors represent different bins sorted by decoded uncertainty within each participant. The black line represents the best linear fit. (I) Correlation between absolute decoding error and decoded uncertainty for individual participants (gray dots). The blue dot represents the group average and the error bar represents ±SEM.

We conducted a planned analysis using a predefined 1-sec time window centered at the middle of the memory delay (1–2 s from delay onset) and computed the power spectral density within this window with Welch’s method (see Methods). Applying the Bayesian decoder to the alpha power (8–13 Hz), we found that the spatial location (polar angle) held in working memory was decodable, as indicated by the absolute decoding errors (Figure 2B and 2C). For the majority of participants (12 out of 14) decoding errors exhibit clustering around 0°, indicating above-chance decoding (Figure 2B for an example subject; see individual results in Supplementary Figure 1). Moreover, the (signed) decoding error significantly correlated with (signed) memory error (Figure 2D and 2E), suggesting a close correspondence between the memory representations in alpha-band activity and behavioral memory reports.

We next investigated whether neural uncertainty of working memory predicts explicit behavioral uncertainty reports. Behavioral uncertainty reports (arc length) increased with the uncertainty derived from the decoded memory content from alpha band (Figure 2F and 2G). This relationship indicates that the memory representations held in alpha-band activity not only supports memory reports, its uncertainty can inform trialwise behavioral uncertainty reports. In addition, we found that the decoded uncertainty correlated with decoding performance, indexed by the (absolute) decoding error, a signature for a well-behaved probabilistic decoder (Figure 2H and 2I).

We applied the decoding analyses to theta (2–7 Hz) and beta power (14–30 Hz) within the same time window. No reliable decoding was observed in the theta band. In the beta band, spatial content was decodable, but the performance was worse than that observed in alpha (Supplementary Figure 2A). In addition, beta-band decoding error did not correlate with behavioral memory error, nor did its decoded uncertainty correlate with reported uncertainty (Supplementary Figures 2B-C).

### Beta-band activity encodes a scalar representation of uncertainty

While the precision of mnemonic representations provides a trial-specific and momentary source of uncertainty, it may not be the sole neural substrate supporting metacognitive judgments. Representational precision, as well as other cues, could be integrated and converted into a simpler format, representing uncertainty or confidence as a scalar, or a magnitude variable. This notion follows the previous findings that regional activity levels or neural population response scale with confidence rating, or opt-out behaviors, in perceptual decisions (e.g., Bang and Fleming 2018; Zylberberg and Shadlen 2025; Fleming et al. 2012; Kiani and Shadlen 2009; Vivar-Lazo and Fetsch 2026). To assess the neural basis of a scalar representation of uncertainty, we applied ridge regression to predict trial-by-trial uncertainty reports (arc length) directly from multivariate band-limited power within the same time window (1–2 s from delay onset) during the memory delay. We found that beta-band activity reliably predicted the uncertainty report, whereas other frequency bands showed chance-level performance (Figure 3).

**Figure 3.**
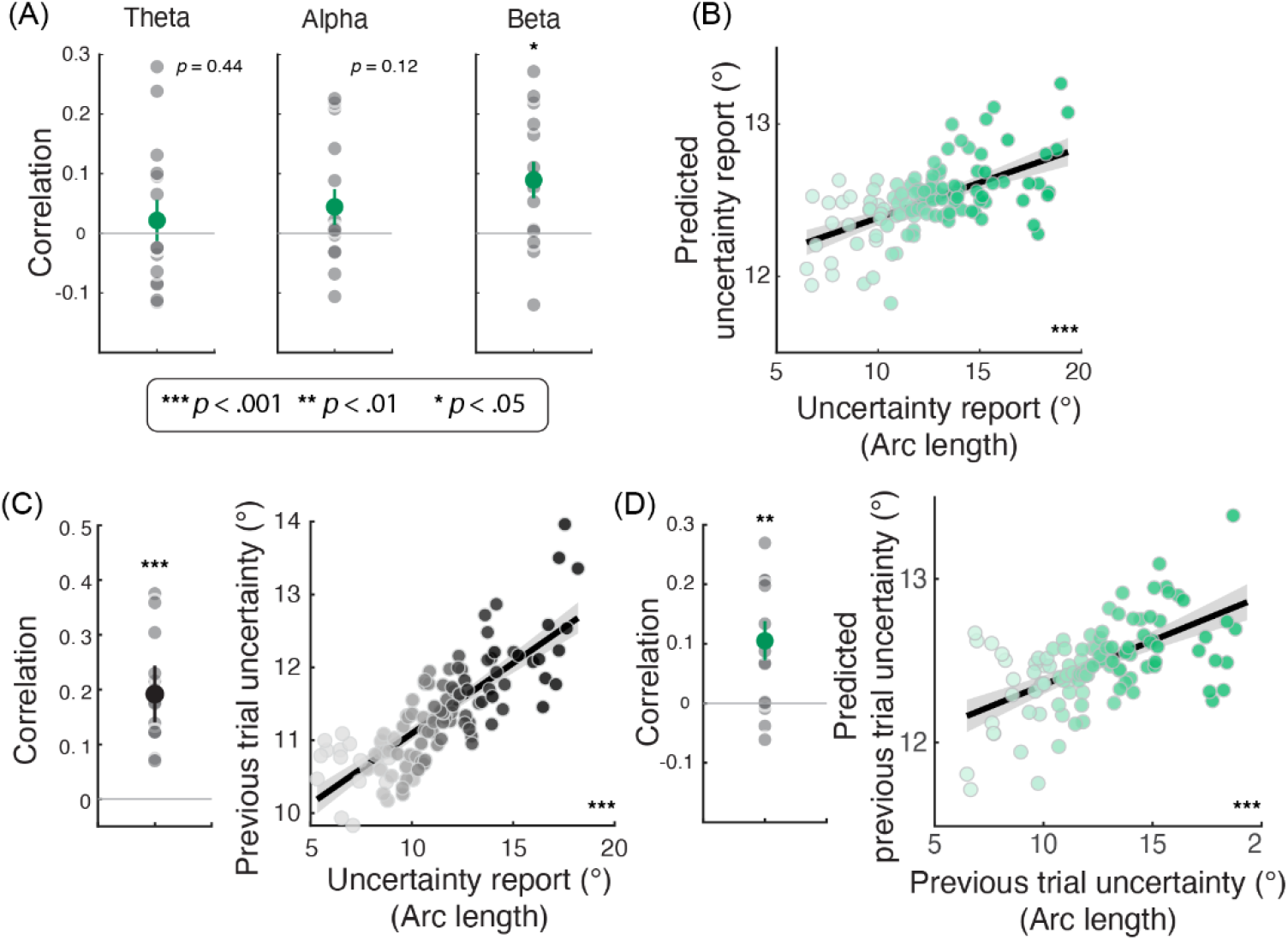
Scalar representation of uncertainty from beta-band activity. (A) Correlations between the uncertainty report and those predicted by ridge regressions using the power of different frequency bands. The gray dots are correlation coefficients for individual participants. The green dot indicates the group mean; error bars denote ±SEM. (B) Predicted uncertainty report from ridge regressions using beta-band activity plotted against the uncertainty report. The black line shows the best linear fit. (C) Serial dependence of uncertainty reports. Correlation between uncertainty reports on the current trial and the previous trial. Data are plotted in the same format as panels (A) and (B). (D) Correlation between uncertainty reports on the previous trial and those predicted by ridge regression using beta-band activity from the current trial. Data are plotted in the same format as panels (A) and (B).

### Distinct time courses and cortical origins of two uncertainty representations

To compare the two forms of uncertainty signals we observed—the representational precision derived by decoding memorized locations from the alpha band, and the scalar uncertainty signal observed in the beta band—we conducted time-resolved decoding and source reconstruction. The time course of decoded uncertainty (from the alpha band) quickly decreased after stimulus onset, indicating decodable information for the target location (Figure 4A). The correlation between decoded uncertainty and behavioral uncertainty reports only emerged during the middle of the delay in a relatively short time window (Figure 4B). In contrast, beta-band activity predicted the reported arc length throughout the entire trial, even before stimulus onset (Figure 4D). This pattern indicates the presence of a persistent, slowly evolving signal that reflects a confidence signal across trials and task epochs. Consistent with this interpretation, behavioral uncertainty reports exhibited serial dependence: uncertainty on the current trial was correlated with that on the previous trial (Figure 3C). To test whether beta-band activity carries information about past uncertainty, we trained a ridge regression model to “predict” the previous trial’s reported arc length from current-trial beta-band activity. Beta-band activity reliably predicted uncertainty from the previous trial (Figure 3D).

**Figure 4.**
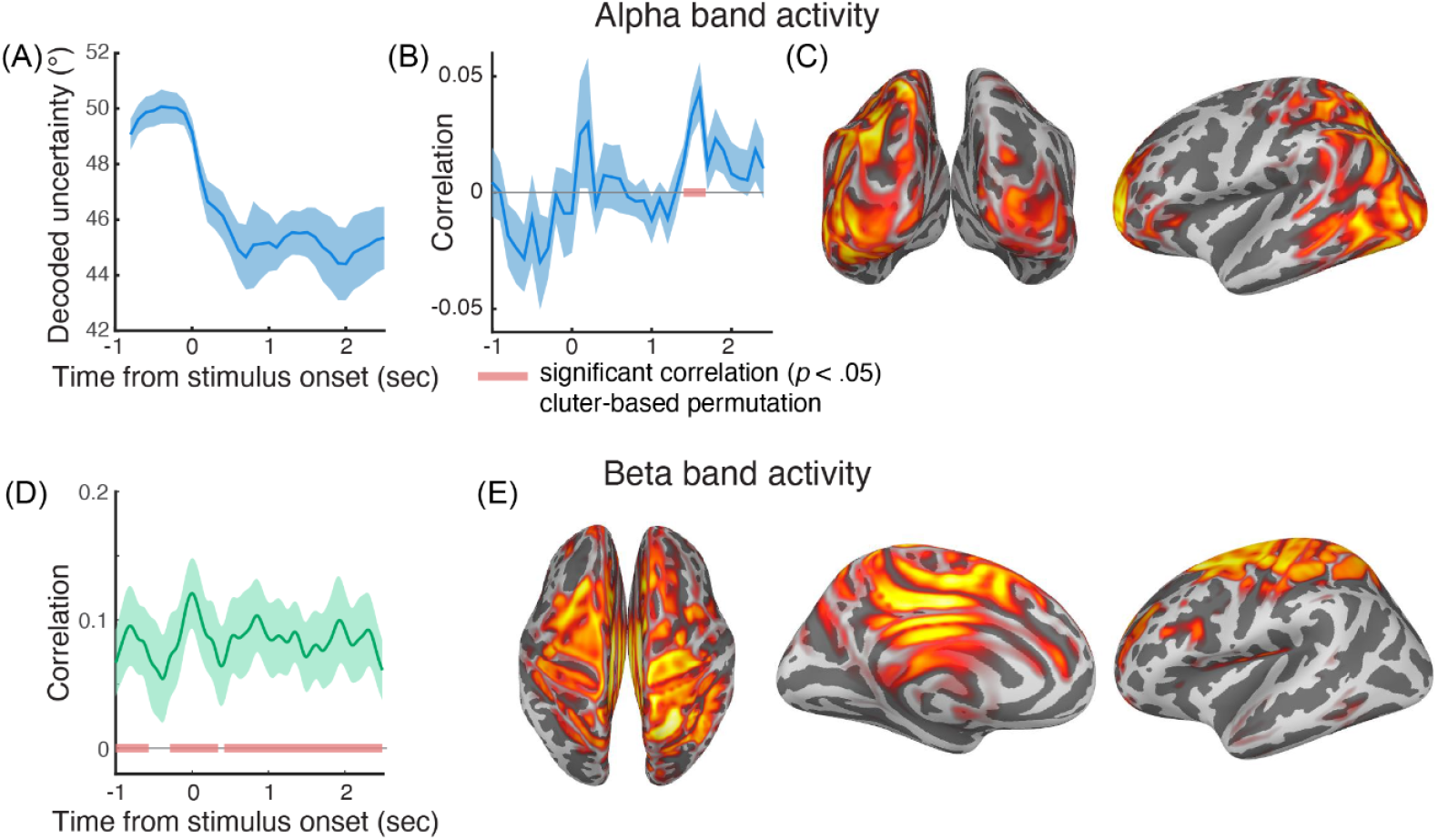
Time-resolved analysis and source reconstructions for two types of uncertainty signals. (A) Time-resolved decoded uncertainty from alpha activity. The blue line represents the group average and blue shaded interval shows ±SEM (B) Correlation between decoded uncertainty and reported uncertainty (arc length). The red line indicates a significant cluster. (C) Source reconstruction for alpha-band activity (power) during the 1-2 s time window. The color represents normalized density (see Source Localization in Methods). (D) Correlation between the predicted current-trial uncertainty report by the beta-band activity, and the actual uncertainty report. The red lines indicate significant clusters. (E) Source reconstruction for beta-band activity (power) during the 1-2 s time window. The color represents normalized density.

We conducted source reconstruction based on the band-limited power during the 1-sec time window in the middle of the memory delay (1 to 2-sec from stimulus onset). The reconstruction showed distinct cortical origins for alpha- and beta-band signals. Alpha-band activity was localized primarily to occipital and posterior parietal regions, including areas along the intraparietal sulcus (Figure 4C). This is consistent with prior work linking alpha oscillations to visual and visuospatial attention brain areas (Capilla et al. 2014; Sokoliuk et al. 2019; Yang et al. 2024). Beta-band activity was instead localized primarily around the central sulcus, encompassing the precentral and postcentral gyri and adjacent sulcal regions (Figure. 4E). The localization of the beta-based signals to sensorimotor cortex suggests that they reflect decision- or action-related content, rather than mnemonic representations per se.

## Discussion

Metacognitive evaluation of one’s own memory is critical for guiding planning and decisions based on information stored in working memory. However, the neural processes that support metacognitive access to working memory have only recently begun to be investigated. Here, we show that working memory uncertainty is expressed in multiple forms in oscillatory neural activity. We identified two distinct but coexisting oscillatory rhythms that convey different aspects of memory uncertainty. Alpha-band activity represents the memorized location and concurrently carries probabilistic information, such that the precision of the mnemonic representation can guide subsequent uncertainty reports. In parallel, beta-band activity carries a scalar representation of uncertainty that tracks a slow-evolving confidence level across task epochs and trials.

Alpha-band activity has been widely linked to spatially specific representations in working memory and attention. In particular, the topography of alpha power can be used to decode the remembered location during delay periods (Foster et al. 2016, 2017; Bae and Luck 2018; Bae 2021; Bender et al. 2025), or the locus of spatial attention (Samaha et al. 2016). Extending this literature beyond decoding the target identity, we show that trial-by-trial decoding error predicts subsequent behavioral error, demonstrating a strong link between alpha-band activity and memory-guided behaviors. More importantly, the precision of the memory representations expressed in alpha-band activity is predictive of subjective uncertainty reports, linking alpha-band activity to metacognitive evaluation of memory quality.

Our recent studies using fMRI has shown that working memory representations reconstructed from BOLD activity in visual and parietal cortex predict both memory reports and uncertainty (Li et al. 2021, 2025), consistent with the idea of a population code that jointly represents an encoded item and its uncertainty (Ma et al. 2006; Jazayeri and Movshon 2006; Walker et al. 2020). Working memory representations are distributed across multiple cortical areas and can be measured through several levels of neural responses (Christophel et al. 2017; Panichello and Buschman 2021; van Kerkoerle et al. 2017). The present results demonstrate that uncertainty-related probabilistic information is also expressed in alpha-band activity, indicating that population codes supporting working memory can be accessed across measurement modalities and temporal scales. Alpha-band signals recorded with EEG reflect large-scale synaptic and transmembrane population dynamics, closely related to local field potentials, and have been widely interpreted as inhibitory modulation that shifts cortical excitability (Jensen and Mazaheri 2010; Mazaheri and Jensen 2010; Iemi et al. 2016; Lange et al. 2013). Consistent with this view, prior work has shown that alpha power is inversely related to BOLD responses during stimulus encoding, suggesting that alpha power and BOLD responses provide complementary information about human brain activity (Hermes et al. 2017). Together with our previous work measuring BOLD responses (Li et al. 2021, 2025), the present findings indicate that this relationship, between alpha power and BOLD responses, extends beyond perception to the maintenance of working memory, and both track the content and uncertainty of mnemonic representations concurrently, indicating a probabilistic population code expressed across multiple measurement scales.

During metacognitive introspection, people rely on multiple sources of information beyond the instantaneous precision of the encoded representation. In our study, we observed a serial dependency in uncertainty reports, whereby the reported uncertainty on one trial correlated with that of the previous trial. Similar history effects or a more global evaluation of self performance have been reported in confidence judgments during categorical perceptual decisions (e.g., Rahnev et al. 2015; Rouault et al. 2019). Consistently, our findings point to the existence of an internal variable carried in the beta-band activity that represents uncertainty as a scalar magnitude variable, which persisted across different trial epochs and trials. This property is well suited for supporting the many behavioral roles attributed to confidence or uncertainty. Confidence has been shown to influence cognitive processing at multiple stages of decision making, as early as how sensory evidence is accumulated (Rollwage et al. 2020), and as late as the execution of motor responses and behavioral reports (Seideman et al. 2018). Post-decision or post-feedback confidence is also known to affect decisions in the subsequent trial (Urai et al. 2017). The slow temporal dynamics of the uncertainty representations in beta-band activity we observed is well-suited to serve such a role, which can continuously influence various phases within and across trials. The scalar representation of uncertainty observed here may not be specific to visual working memory. A recent study on probability learning also reported a persistent confidence signal carried by beta-band activity, even during the pre-trial period, suggesting that beta-band dynamics may reflect a more general confidence state (Meyniel 2020). While there are differences in the nature of these signals, our results are broadly in line with recent findings from non-human primate neurophysiology, reporting that pre-trial baseline activity in the dorsolateral prefrontal cortex (DLPFC) neural population is indicative of subsequent metacognitive judgments of working memory (Ning et al. 2026).

The localization of this beta-band activity to regions around the pre- and post-central sulcus is in line with previous work showing that beta-band activity in sensorimotor cortex can index a derived confidence estimate of an ideal observe model in motor-adaptation tasks (Tan et al. 2016), and with evidence that TMS disruption of premotor cortex reduces serial dependence in visual estimation tasks, implicating these areas’ contribution in maintaining information from one trial to the next (de Azevedo Neto and Bartels 2021).

Earlier work has proposed that beta oscillations reflect the maintenance of the “status quo,” where beta-band activity is associated with stabilizing the current motor state and suppressing changes in action plans. Related notions have been extended to various cognitive domains, suggesting that beta-band activity may support the stabilization of ongoing cognitive states, or to serve various top-down control functions in visual attention and working memory tasks (Kilavik et al. 2013; Engel and Fries 2010; Fries 2015; Lundqvist et al. 2018; Liljefors et al. 2024; Buschman et al. 2012). The other account on beta-band activity has emphasized that beta oscillations also represent content-specific information, such as the parametric features of the items held in working memory (Spitzer and Haegens 2017; Wimmer et al. 2016; Spitzer and Blankenburg 2011). Our findings reflect that these different interpretations on beta-band activity may be complementary. In our task, beta-band activity can be viewed as maintaining an internal state of uncertainty or confidence that persists across time. Importantly, the notion of ‘state’ here is not a distinct mode devoid of content. Rather, like many latent variables in computational models of decision-making, the state carries a specific numerical value, much like the content maintained in working memory (e.g., the color or orientation of an object). In the present case, the maintained value is the magnitude of uncertainty itself.

## Methods

### Participants

In this study, we recruited 14 participants (7 males; 22–28 years old). All participants were right-handed and had normal or corrected-to-normal vision. They provided written informed consent prior to the experiment. The study was approved by the Ethics Committee of Tianjin University in accordance with institutional guidelines.

### Procedures

The experimental procedure was implemented using Psychophysics Toolbox Version 3 running on MATLAB R2017a. Each participant completed 20 runs, with each run consisting of 32 trials. The trials were separated by a 2-3 s inter-trial interval, during which the fixation point was in dark gray. At the beginning of each trial, the fixation point increased in brightness to indicate trial onset. Participants were required to maintain their gaze at the fixation point for 1 s. The target, a bright dot, was then presented for 500 ms. The eccentricity of the target was fixed at 10° visual angle, and its polar angle was pseudo-randomly selected from 32 equally spaced positions spanning the full 360° range. After target offset, participants continued to fixate for a 2 s memory delay period while minimizing blinks and eye movements.

At the end of the delay, a black ring appeared on the screen as a response cue. Participants first moved a cursor to report the remembered target location and could fine-tune their response using the Up and Down arrow keys on the keyboard. After finalizing their memory report by pressing the space bar on the keyboard, an arc with a random initial length appears on the screen, centered at the reported target location with equal extension on both sides. Participants adjusted the length of the arc using the Up and Down arrow keys on the keyboard with both sides of the arc changing simultaneously. They pressed the space key again to confirm the reported arc length. Finally, a feedback dot was presented at the true target location. Reward points were determined by the reported arc length and whether the true target position fell within the arc.

The reward was designed to incentivize well-calibrated uncertainty reports (Li et al. 2021). The number of points (100^*e*−0.08*d*^, where *d* is the length of the arc) participants could earn decreased monotonically with the length of the reported arc, such that shorter arcs yielded higher potential rewards. However, points were awarded only if the true target location fell within the reported arc; otherwise, participants earned zero points. This created a trade-off between maximizing potential reward and ensuring sufficient coverage to include the target. The optimal strategy was to adjust the arc length according to one’s subjective uncertainty or confidence in the memory report (Yoo et al. 2018; Honig et al. 2020).

### EEG setup

EEG data were recorded using a Neuroscan SynAmps2 system with 62 Ag/AgCl electrodes placed according to the international 10–20 system. The EEG signals were sampled at 1000Hz, and electrode impedances were maintained below 5 kΩ throughout the recording. During each experiment, eye movements and pupil diameter were recorded using a Tobii eye tracker (Tobii Pro Spectrum) at a sampling rate of 600 Hz.

### EEG preprocessing

EEG data were preprocessed using the EEGLAB toolbox (Delorme and Makeig 2004) in MATLAB. The data were re-referenced to the average reference, notch-filtered at 50 Hz and band-pass filtered between 0.1 and 45 Hz, and resampled to 500 Hz. Bad trials were identified and removed for all participants, followed by independent component analysis (ICA) to remove ocular, muscle, and other stereotypical artifacts.

We focused primarily on the 1 s time window during the memory delay period (1–2 s after target onset). Subsequent time-course and time-frequency analyses (Figure 4) were based on epochs ranging from −1000 ms before to 2500 ms after stimulus onset. In total, 8,960 trials were collected from 14 participants (640 trials each). After bad-trial exclusion, 8,546 trials were retained, and approximately 4% were excluded.

Feature extraction was performed using the MNE-Python package (Gramfort et al. 2013). A random signal typically has finite average power, allowing it to be described in terms of its average power spectral density (PSD) *S*_*x*_ (*f*), which is defined as:

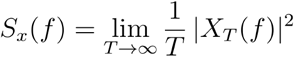

Where *X*_*T*_ (*f*) is the Fourier transform of a finite time window, and *T* is the length. Here, the fixed-window (1-2 s from target onset) PSD for different frequency bands was estimated using Welch’s method with a Hamming window and FFT length was set to 256.

Time-frequency representation of the EEG signals were computed using the Short-Time Fourier Transform (STFT). Unlike the classical Fourier transform, which provides only global frequency information, STFT allows the spectral content of a signal to be examined as it evolves over time by applying the Fourier transform within short temporal windows.

Given a signal *x*(*t*), the STFT is computed by multiplying the signal with a sliding window function *w*(*t*) centered at time *τ*, and then performing a Fourier transform within that window:

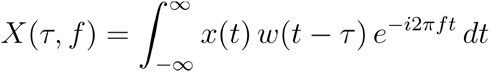

Where *w* (*t* − *τ*) is window function centered at time *τ* and *f* is frequency. In this study, a Hann window was applied and window length was set to 500 ms. The Fourier transform was computed with a sliding window to estimate spectral power over time.

### EEG Decoding

#### Bayesian decoding

We conducted Bayesian decoding using the TAFKAP algorithm (van Bergen and Jehee 2021; van Bergen et al. 2015) to perform Bayesian decoding, extracting working memory representations and quantifying their uncertainty in probabilistic terms. This method allowed us to read out both the remembered location and the associated memory uncertainty jointly for each single trial.

In the generative model, the multivariate EEG signal associated with a target location (polar angle) was modeled as a multivariate normal distribution. The expected response (mean) of each electrode was defined by its tuning curve, representing an electrode’s response (e.g., alpha power) as a function of target polar angle.

The tuning function of each electrode *f*(*s*) was modeled as a linear combination of eight basis functions *g*_*k*_(*s*), which evenly spaced the polar angle space. These basis functions were raised cosine functions

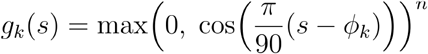

where *ϕ*_*k*_is the center of *k*th basis. The activity of *i*-th electrode to given the stimulus *s* is modeled as

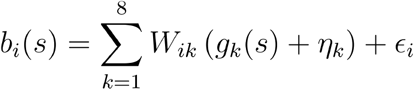

where *W*_*ik*_ weights the basis function’s contribution to the *i*-th channel’s tuning curve. The η and ∈ are the basis-specific noise and electrode-specific noise respectively. The probability of measuring EEG activity pattern **b** given the stimulus is then *p*(**b** ∣*s; θ*) = *N*(*f*(*s*), Ω). Where the model parameters *θ* are the basis function weights **W** and covariance matrix Ω. Here, the covariance is modeled as Ω = *λ*Ω_0_ + (1 - *λ*) Ω_sample_.

To ensure a stable estimation of the covariance matrix, we modeled the covariance as a shrinkage of the sample covariance Ω_sample_ toward a target covariance matrix Ω_0_. The free parameter *λ*determined the degree of shrinkage. Here, the target covariance matrix has a simple structure, computed as a weighted sum over a diagonal matrix, a rank-1 covariance matrix, and the covariance depending on the tuning weight matrix of the voxels (see details in (van Bergen and Jehee 2021; van Bergen et al. 2015)).

We estimated the free parameters by maximizing the likelihood of the training data. We then decoded the target location on each trial in the test set based on the EEG activity pattern using Bayesian inference. Specifically, the posterior distribution was computed via Bayes’ rule.

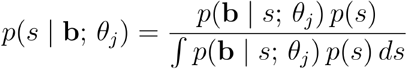

We defined the mean and standard deviation of the trialwise posterior distribution as the decoded location and decoded uncertainty, respectively. We adopted the bagging procedure from the TAFKAP algorithm, in which the training data were bootstrapped to decode trials in the test set repeatedly, and the resulting posterior distributions were averaged until stable estimates were obtained (van Bergen and Jehee 2021).

#### Ridge regression

To decode the neural activity that underlies scalar representations of uncertainty we used ridge regression to directly predict the report uncertainty (arc length) by the power of different frequency bands. Ridge regression is a regularized linear regression method by adding an *L*_2_ penalty term to ordinary least squares (OLS), which shrinks the magnitude of the coefficients.

Given the arc length *d* and the EEG activity *B*, ridge regression estimates the coefficient vector *w* by minimizing

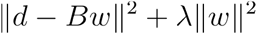

where *λ* is a non-negative regularization parameter controlling the degree of shrinkage applied to the regression coefficients. In our analysis, the regularization parameter was set to *λ =* 10.

Here, we used a leave-one-run-out cross-validation scheme, where run index according to the run order in the experiment. The data contain 20 runs, we trained the model on 19 runs to obtain the regression coefficients, and then used these coefficients to estimate *d* on the remaining test run.

#### Source Localization

We used Python-MNE to perform the source localization and visualization. sLORETA was used in Figure 4C and 4E to estimate cortical current density from EEG activity (Pascual-Marqui 2002). It belongs to the family of minimum-norm (MNE) inverse solutions.

The FreeSurfer fsaverage template was used to construct the source space (spacing = ‘oct6’). The boundary element model (BEM), and the forward model were estimated for subsequent source reconstruction. The noise covariance matrix was estimated from the baseline interval (−0.5 to −0.05 s before the target onset). EEG signals were filtered at the frequency band of interest, averaged across trials within the time window of interest (1–2 s from target onset) before performing source reconstruction.

Cortical sources were estimated by inverting the forward model using sLORETA. The resulting source estimates were originally in arbitrary units and reflect relative current density. In sLORETA, source estimates were further normalized by their expected variance, given by the diagonal of the resolution matrix. This normalization accounts for differences in sensitivity across cortical locations, with variance primarily determined by the forward model geometry (e.g., cortical depth and sensor configuration) and the noise covariance estimated from the baseline interval.

#### Quantification and statistical analysis

In multiple figures, we illustrate the correlations between two variables (Figure 1E; Figure 2E,G,I; Figure 3A). Pearson correlation coefficients were first computed within each participant. The resulting coefficients were Fisher z-transformed, and group-level significance was assessed using a one-sample t-test against zero. Alongside the single-trial correlations, we present scatter plots for visualization purposes. In these plots, each participant contributed eight data points (bins). For each participant, trials were first sorted according to the variable shown on the x-axis and then divided into eight bins with equal number of trials. For each bin, we computed the mean value of the variable on the y-axis across trials within that bin. Before pooling the data across participants, each participant’s values were mean-centered to remove individual differences (for both x- and y-axis). As a result, between-subject variability did not contribute to the scatter plots.

Cluster-based permutation test was applied to test the time-resolved correlations reported in Figure 4. The procedures are based on those developed in (Maris and Oostenveld 2007). For Figure 4B, we computed the correlation between decoded uncertainty (through Bayesian decoding of target location on alpha-band activity) and reported uncertainty within each participant. At each time point, group-level significance was evaluated using a one-sample t-test against zero. Adjacent time points with uncorrected significance (*p* < 0.05) were grouped into clusters, and cluster statistics were defined as the sum of t-values within each cluster. Statistical significance was determined by comparing the observed cluster statistics to a null distribution generated by randomly permuting the reported uncertainty across trials and repeating the analysis. For each permutation, the maximum cluster-level t-sum was recorded. This procedure was repeated 1,000 times to form the null distribution, and clusters exceeding the 95th percentile were considered significant. The same procedure was applied in Figure 4D to test the correlation between predicted uncertainty (from ridge regression on beta-band activity) and reported uncertainty.

---

**Supplementary Figure 1.**
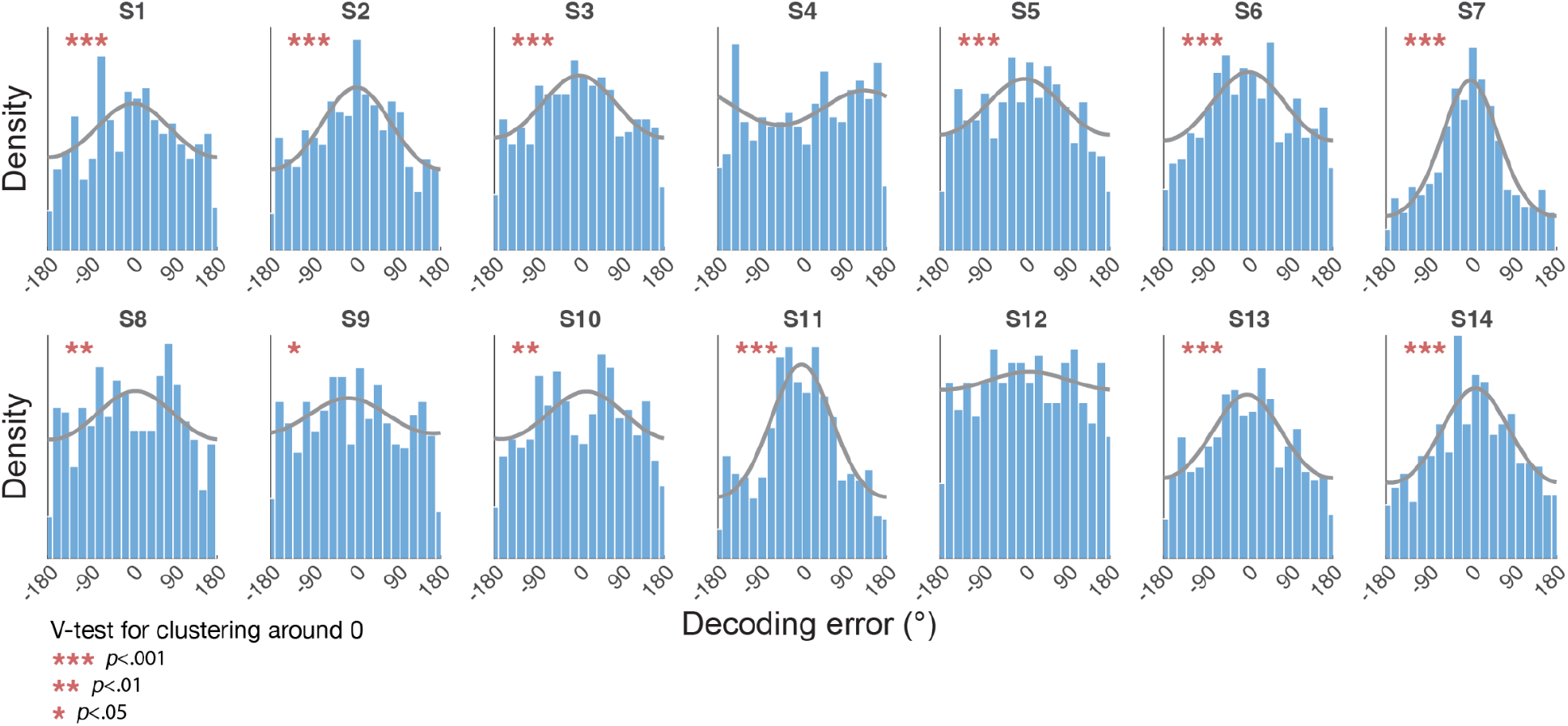
Distribution of decoding errors for individual participants. Histograms show decoding errors from alpha-band activity. Gray lines indicate best-fitting von Mises distributions. Decodability was assessed for each participant using a circular V-test on the error distribution. Significant V-tests indicate non-uniformity with clustering around 0°. Twelve of fourteen participants showed significant decoding.

**Supplementary Figure 2.**
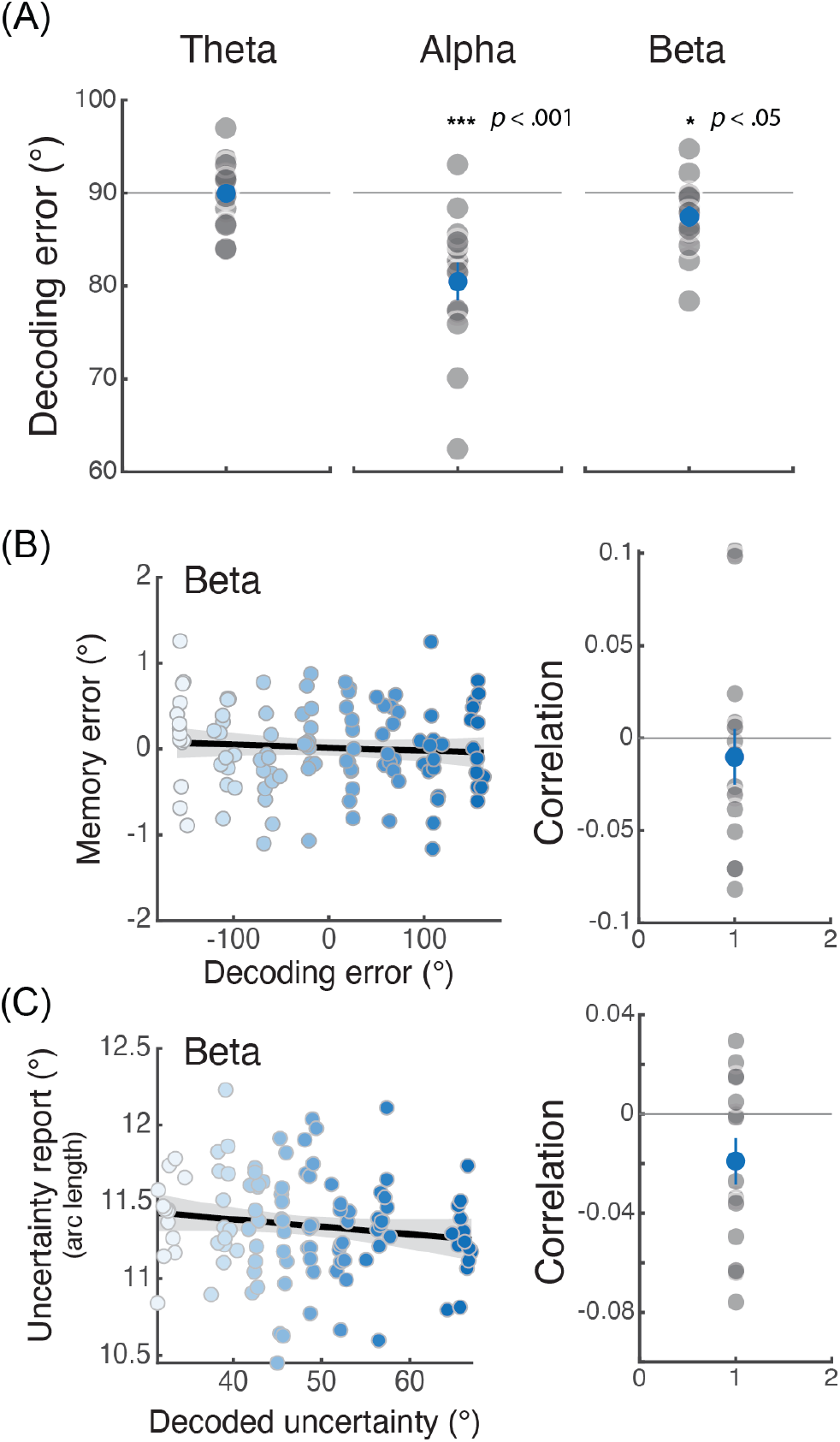
(A) Absolute decoding error from theta, alpha and beta power. The results of alpha are those plotted in Figure 2C. Gray dots represent individual participants. The blue dot represents mean ±SEM. Distribution of decoding errors for individual participants. (B) Correlations between decoding error and memory error based on decoding of beta power, plotted in the same format as in Figure 2D and 2E. (C) Correlations between decoding error and memory error based on decoding of beta power, plotted in the same format as in Figure 3.

## Notes

Conflicts of interest: The authors declare no competing financial interests.

### Competing Interest Statement

The authors have declared no competing interest.

